# DNA replication errors drive genome-wide small inverted triplication dynamics

**DOI:** 10.64898/2025.12.26.696596

**Authors:** Yi Lei, Yu Zhou, Haitao Sun, Hang Yuan, Xinyu Pei, Jessica D. Hess, Yao Yan, Zunsong Hu, Mian Zhou, Zhaohui Gu, Li Zheng, Xiwei Wu, Binghui Shen

## Abstract

Structural variants (SVs) have a profound impact on phenotype and diversity and are associated with human diseases. To explore the origination of SVs, we have analyzed 1,340 cancer genomes with annotation of 4,608 novel small inverted triplication (SIT) events and found that *FEN1* is strongly associated with SIT incidence. Then, we performed long-read sequencing and developed PacBioR to annotate SITs in yeast *FEN1* mutant cells. We found that SIT structures mimic classic inverted triplications but with a smaller DUP/IN/DUP structure of 184/160/184 bp on average, with a spacer sequence of 30 bp and breakpoint junction of 6 bp. We further showed that breakpoints of SITs preferentially occurred at nucleosome midpoints, aligned with Okazaki fragment termini, and those harbored in plasmids were precisely eliminated via DNA polymerase slippage over SIT-derived hairpin structures. This study provides mechanistic insight into SIT origination and offers practical tools for future studies on genome rearrangements.

## 1. Introduction

Genome stability is essential for cell survival and normal growth but can be compromised by structural variants (SVs) such as insertions, deletions, inversions, and duplications.^[1–3]^ Depending on the genomic context and affected genes, SVs are associated with many human diseases including, but not limited to, cancers.^[4]^ One of the most important endeavors of the biomedical community is the discovery and characterization of SVs across human genomes.^[5]^

Conventional technologies such as karyotyping,^[6]^ fluorescence in situ hybridization,^[7,8]^ and comparative genomic hybridization arrays^[9]^ have served as important methods for detecting and validating SVs. However, their limited sensitivity and resolution constrain the genome wide detection in proliferating cell populations and hinder the discovery of novel SVs. High-throughput sequencing technologies have now become indispensable for investigating SVs at the whole-genome level. Various short-read-based approaches for SV detection have been developed, which typically rely on strategies such as read-depth analysis,^[10]^ discordant read-pair mapping,^[11]^ split-read alignment,^[12]^ local assembly,^[13]^ or combinations of these methods.^[14–16]^ Although short-read technologies have been widely applied in large-scale projects, including the 1000 Genomes Project,^[17,18]^ they are not ideal to detect and characterize SVs. In contrast, long-read sequencing provides several advantages for SV detection.^[19–22]^ In particular, PacBio high-fidelity (HiFi) sequencing generates long (10–25 Kbp) and highly accurate (> 99.9%) sequencing reads,^[23]^ making it well-suited for characterizing SVs.^[24,25]^ However, until the current work, the identification of SVs has been challenging,^[26]^ particularly for rare and complex SVs.

One specific type of SVs, the inverted triplication, consists of an inverted central copy flanked by two direct copies (DUP/IN/DUP),^[27]^ and was first identified using fluorescence in situ hybridization.^[28]^ Since then, increasing numbers of locus-specific inverted triplications have been reported, and accumulating evidence indicates that such structures can occur on nearly every human chromosome.^[27,29–39]^ To date, studies of individual inverted triplication cases have led to three proposed models for their formation.^[40,41]^ Brewer et al. observed that an inverted triplication is generated at the *SUL1* locus when the budding yeast *Saccharomyces cerevisiae* is grown long-term in sulfate-limiting chemostats and proposed the origin-dependent inverted-repeat amplification (ODIRA) model.^[41]^ This model suggests that replication errors at pre-existing, interrupted inverted repeats can generate extrachromosomal inverted dimeric intermediates, which can autonomously replicate and reintegrate into the genome, thereby forming an inverted triplication.^[27,41,42]^ Carvalho et al. analyzed a disease-specific cohort with an inverted triplication that occurred at the *MECP2* locus and proposed that these structures can arise through a combination of homology-directed break-induced replication (BIR) and template switching, or through microhomology-mediated BIR and nonhomologous end joining.^[40]^ In a subsequent study, they proposed an iterative template switching model, in which replication forks encountering inverted sequences undergo multiple rounds of template switching, ultimately producing an inverted triplication.^[29]^

In the current study, we report a novel SV that shares structural features with the identified inverted triplications but is smaller in size than those previously described,^[27,41,42]^ with a median size of 184/160/184 bp (DUP/IN/DUP), which we characterize as a small inverted triplication (SIT) arising from DNA replication errors. Using 1,340 short-read sequencing datasets, we identified the occurrence of SIT in different cancer genomes. Subsequent mutational analysis suggested that 4,002 genes with functional deficient mutations were significantly enriched in the high-incidence SIT group, and *FEN1* was among the top 10 most significantly enriched genes. We performed long-read sequencing of the *FEN1* yeast homologue (*rad27Δ*) mutant and 11 other strains, and developed PacBioR to detect SVs, including SITs, in the HiFi sequencing data. The results showed a marked increase in the incidence of SIT events in the *rad27Δ* mutant compared to the other strains. The breakpoints of SITs preferentially occurred at nucleosome midpoints and aligned with Okazaki fragment termini. Moreover, we found that SIT structures are inherently unstable within cells, and that knockout of *sbcC*, *sbcD*, or *rep* suppressed their elimination, whereas deletion of *dnaQ* or *holC* promoted such a loss. Taken together, our study provides the most comprehensive insights to date into the formation and elimination mechanisms of SITs.

## 2. Results

### 2.1. *FEN1* mutations exhibited a striking enrichment in the high SIT incidence cancer population

We first characterized SIT structures in B-ALL patients (**Figure S1** and **Table S1**). To determine whether such structure could be detected in other cancers, we analyzed 1,340 short-read sequencing datasets from patients representing 22 cancer types and identified 4,608 SIT events (**Figure 1A**, **Table S1**, **Table S2**, and **Figure S1B-C**). To explore the genetic basis underlying SIT formation, we performed a mutational analysis and identified 17,727 genes with functional deficient mutations. To evaluate the association of SIT events with gene mutations, we defined two groups: Samples with ≥ 20 SIT events, corresponding to 90th percentile of non-zero cases, as group 1 (*n* = 60) and with zero events as group 2 (*n* = 745) (**Figure 1B**, **Table S1**, and **Table S2**). Comparative analysis showed that a total of 4,309 genes were significantly enriched (Odds ratio (OR) > 2 and FDR < 0.05), including 4,002 and 307 genes in groups 1 and 2, respectively (**Figure 1C** and **Table S3**). For the genes significantly enriched in group 1, the Gene Ontology (GO) analysis showed that the enriched mutant genes were involved in nucleic acid metabolism, such as RNA splicing, DNA replication, recombination, and repair (**Figure S2** and **Table S4**). When we focused on the 1,179 DNA-related genes identified by the GO and KEGG databases (**Table S5**), the results showed that 303 and 11 genes were enriched in group 1 and group 2, respectively (**Figure 1C** and **Table S5**). The top 10 genes with the highest OR value are shown in **Figure 1D**. Among them, *FEN1* mutations exhibited a striking enrichment in the high SIT incidence group (**Figure 1D** and **Table S3**).

**Figure 1.**
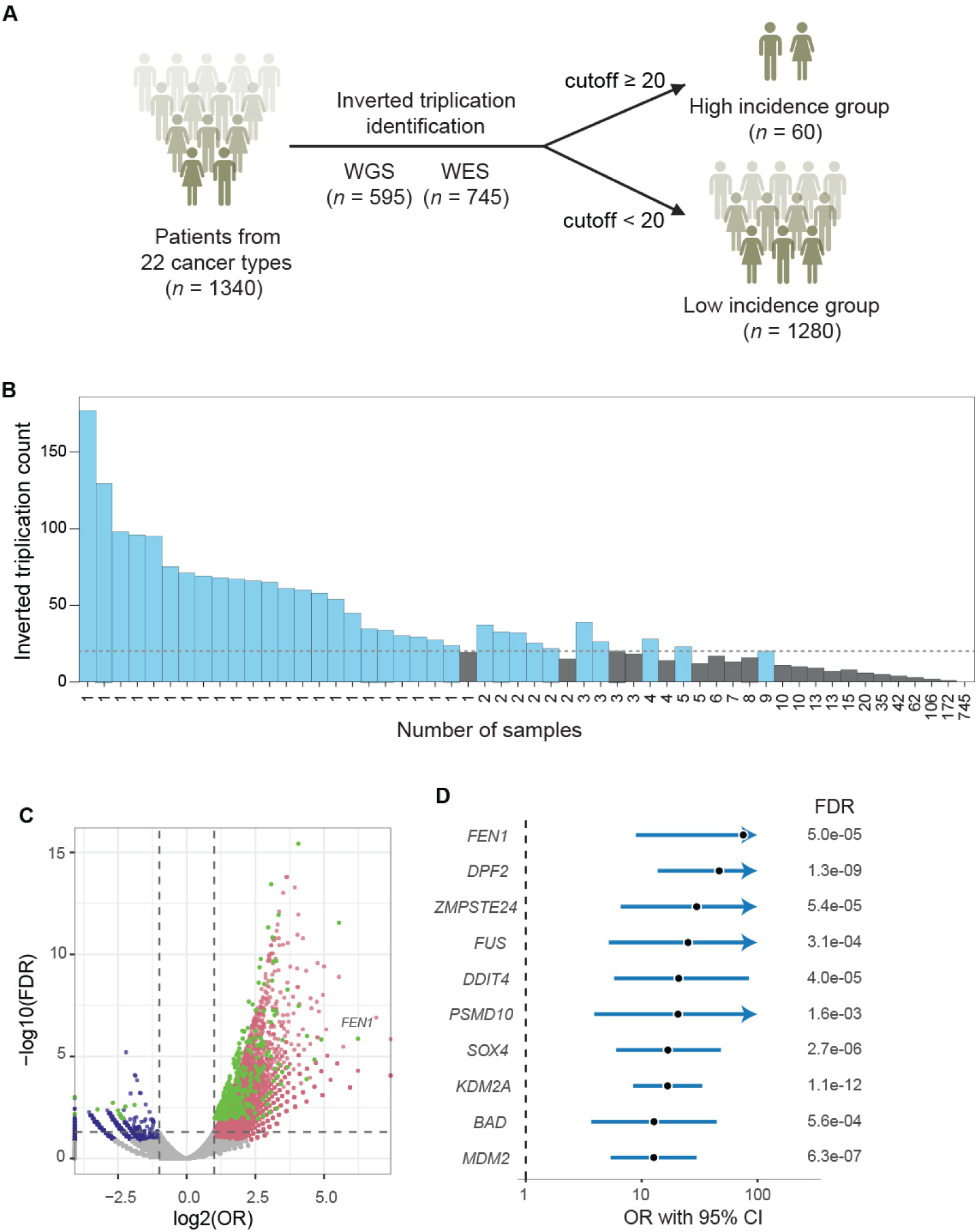
SIT incidence is shaped by individual genetic background. **(A)** A total of 1,340 samples spanning 22 cancer types were analyzed to detect SIT events (**Table S1**). **(B)** Distribution of 4,608 SIT events across individual samples (**Table S1**). Samples were classified into high and low IT incidence groups using a cutoff of 20 SIT events per sample, corresponding to 90th percentile of non-zero cases. **(C)** Volcano plot of all mutated genes, highlighting those enriched in group 1 (log2OR > 1, FDR < 0.05) or in group 2 (log2OR < −1, FDR < 0.05). DNA-related genes, including those involved in DNA replication, repair, and damage response, are highlighted in green. **(D)** The top 10 DNA-related genes with the highest OR values are shown (**Table S3**).

### 2.2. PacBioR: a high-performance tool for accurate annotation of SITs with HiFi sequencing data

Although we confirmed the occurrence of SITs across different cancers, short-read sequencing has inherent limitations to identify complex SVs, as it often fails to span entire SV regions, which may result in incomplete breakpoint information and inaccurate assembly.^[43,44]^ To facilitate a more comprehensive investigation of SIT structure, we developed PacBioR, an R package specifically designed to identify SVs, including SITs, from PacBio HiFi reads (**Figure 2A**). To evaluate the performance of PacBioR, we simulated 2,150 SVs on the hg38 genome using Visor^[45]^ and Sim-it.^[46]^ These SVs contain 1,000 insertions and deletions, and 50 duplications, inversions, and inverted triplications. For insertion and deletions, we simulated them directly using Sim-it to generate PacBio HiFi long reads with 10× and 20× genome coverage (**Figure 2B-C** and **Table S2**). For inversions, duplications, and inverted triplications, we simulated 50 events each using Visor and generated 10× and 20× PacBio HiFi reads with the modified hg38 genome using Sim-it (**Figure 2B-C** and **Table S6**). In addition to PacBioR, we also included other popular SV detection algorithms, including PBSV v.2.11.0 (https://github.com/PacificBiosciences/pbsv), Sniffles v.2.6.3,^[47]^ SVision v1.4,^[24]^ cuteSV v.2.1.2,^[22]^ and SVIM v.2.2.0.^[48]^

**Figure 2.**
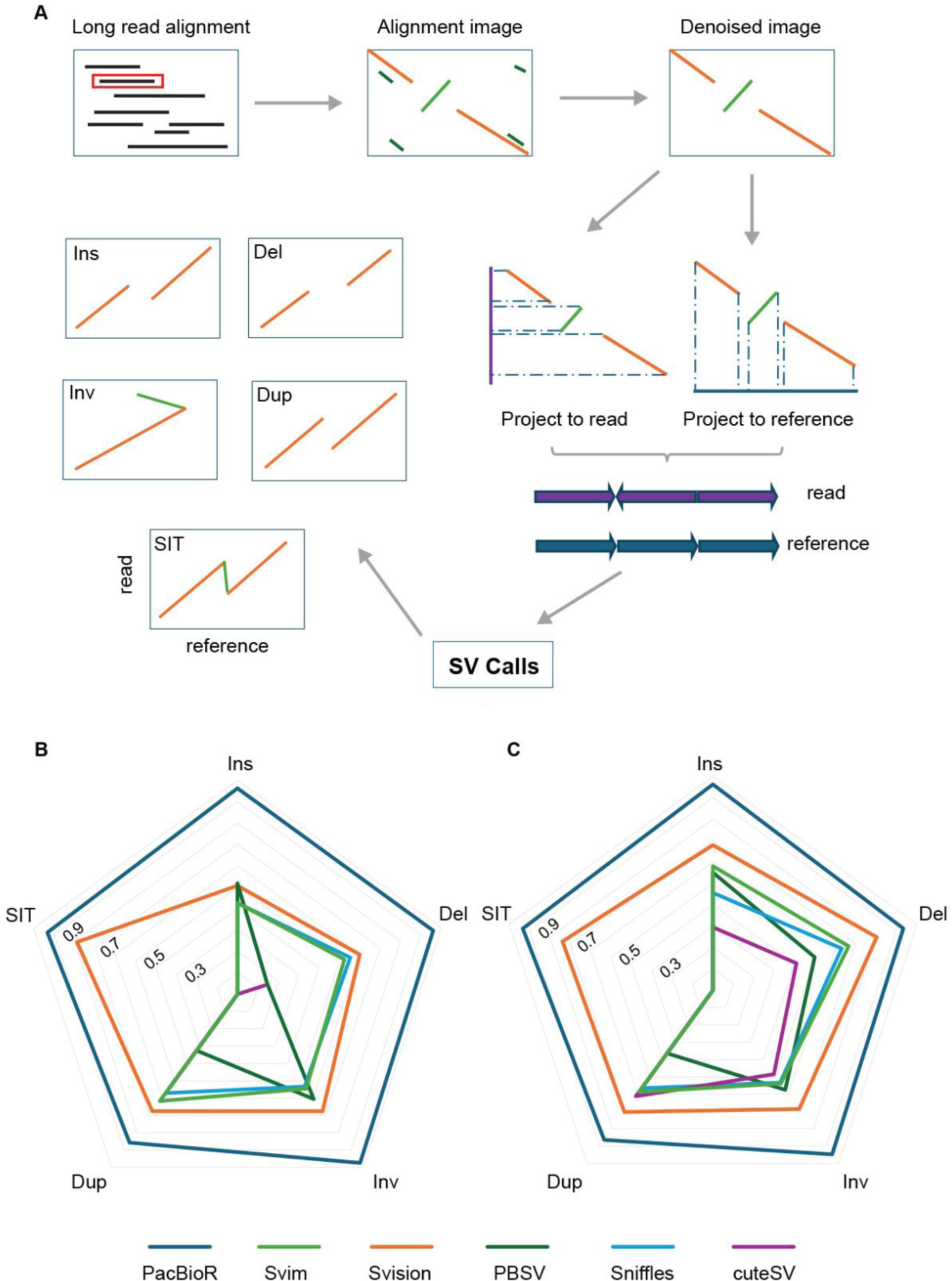
The high-performance PacBioR compared with different SV callers. **(A)** Schematic representation of the PacBioR algorithm. The F1 score of simulated SV detection using different tools is shown in **(B)** and **(C)**, corresponding to **Table S6**. HiFi reads were simulated using a modified hg38 genome with different SVs, with 10x genome coverage **(B)** and 20x genome coverage **(C)**. DEL, INS, INV, DUP, and SIT represent deletion, insertion, inversion, duplication, and small inverted triplication.

PacBioR achieved high precision and recall for all types of SV events, at both coverage levels (**Figure 2B-C** and **Table S2**). Almost all tools, except PBSV, could detect deletion events with high accuracy, with SVision having the highest precision and F1 statistics, and PacBioR having the highest recall for deletions. It is interesting to note that all tools except PacBioR suffered low precision for insertions, suggesting insertions are more difficult to detect. SVision is the only other tool that detected inversions with similar accuracy to PacBioR. Moreover, PacBioR detected duplications with much higher accuracy than all other tools, mainly due to its higher recall rate. As SIT is one type of complex SV, besides PacBioR, only SVision could detect it among the tools we evaluated. While SVision has similar precision as PacBioR, it suffers a low recall rate (60% at both 10× and 20× coverage) (**Figure 2B-C** and **Table S6**). Overall, PacBioR outperformed all other tools when detecting insertions, duplications, inversions, and SITs, while displaying similar accuracy to the other tools for deletion events (**Figure 2B-C** and **Table S6**).

### 2.3. Loss of the *rad27* gene promotes SIT formation

To validate the role of FEN1 in the formation of SITs, we performed HiFi sequencing on wild type (WT) and *rad27Δ* yeast cells (**Table S7**). HiFi sequencing of the WT and *rad27Δ* strains yielded read depths of 919× and 763×, and total reads of 1,102,573 and 1,005,166, respectively (**Table 1** and **Table S7**). Using PacBioR, we identified 742 reads containing SITs in the *rad27Δ* strain, which were clustered into 95 unique events based on their genomic positions, whereas no such events were detected in the WT (**Table 1** and **Table S8**). To validate these results, we performed independent replicate experiments on both strains, which exhibited high reproducibility: WT and *rad27Δ* yielded zero and 95 SITs, respectively (**Table 1** and **Table S8**).

**Table 1.** Comparative analysis of SIT events in 12 yeast strains.

| Sample | Total reads | SIT reads | SIT events | °C |
| --- | --- | --- | --- | --- |
| wild type | 946283 | 0 | 0 | 30 |
|  | 1102573 | 0 | 0 | 30 |
|  | 922423 | 0 | 0 | 37 |
| *rad27Δ* | 913064 | 858 | 95 | 30 |
|  | 1005166 | 742 | 95 | 30 |
|  | 876200 | 986 | 230 | 37 |
| *rad27Δ_dun1Δ* | 858653 | 182 | 35 | 30 |
|  | 772996 | 186 | 17 | 37 |
| *dna2-1* | 952111 | 5 | 5 | 30 |
| *dun1Δ* | 1115480 | 1 | 1 | 30 |
| *exo1Δ* | 959880 | 2 | 2 | 30 |
| *mre11Δ* | 922525 | 0 | 0 | 30 |
| *mus81Δ* | 600256 | 2 | 2 | 30 |
| *rad1Δ* | 984584 | 1 | 1 | 30 |
| *sae2Δ* | 825403 | 1 | 1 | 30 |
| *sgs1Δ* | 999977 | 0 | 0 | 30 |
| *rad51Δ* | 918747 | 0 | 0 | 30 |

Rad27 is a key nuclease that removes the 5’ flap structure during the OFM process.^[49,50]^ Failure to process the 5’ flap may contribute to the formation of SITs. Evidence has been provided that EXO1 and DNA2 also participate in removing 5′ flaps during the OFM process; thus, we generated mutant yeast strains for these two genes (**Table S7**). We detected five SITs in the *exo1Δ* strain, whereas only one was observed in the *dna2Δ* (**Figure 3A** and **Table S8**). Our previous study showed that restrictive temperature (37℃) activates Dun1, which facilitates the transformation of unprocessed 5’ flaps into 3’ flaps, thereby reinitiating DNA extension.^[51]^ To test this, we did HiFi sequencing analysis on WT, *rad27Δ, dun1Δ*, and *rad27Δ dun1Δ* double knockout strains and detected 0, 230, 0, and 35 SIT events, respectively (**Table 1** and **Table S8**). The number of SIT events in *rad27Δ* mutants at 37℃ increased by ∼2.5-fold compared to the normal condition and decreased ∼6.5-fold when the *dun1* gene was knocked out (**Table 1**, **Table S7**, and **Table S8**).

**Figure 3.**
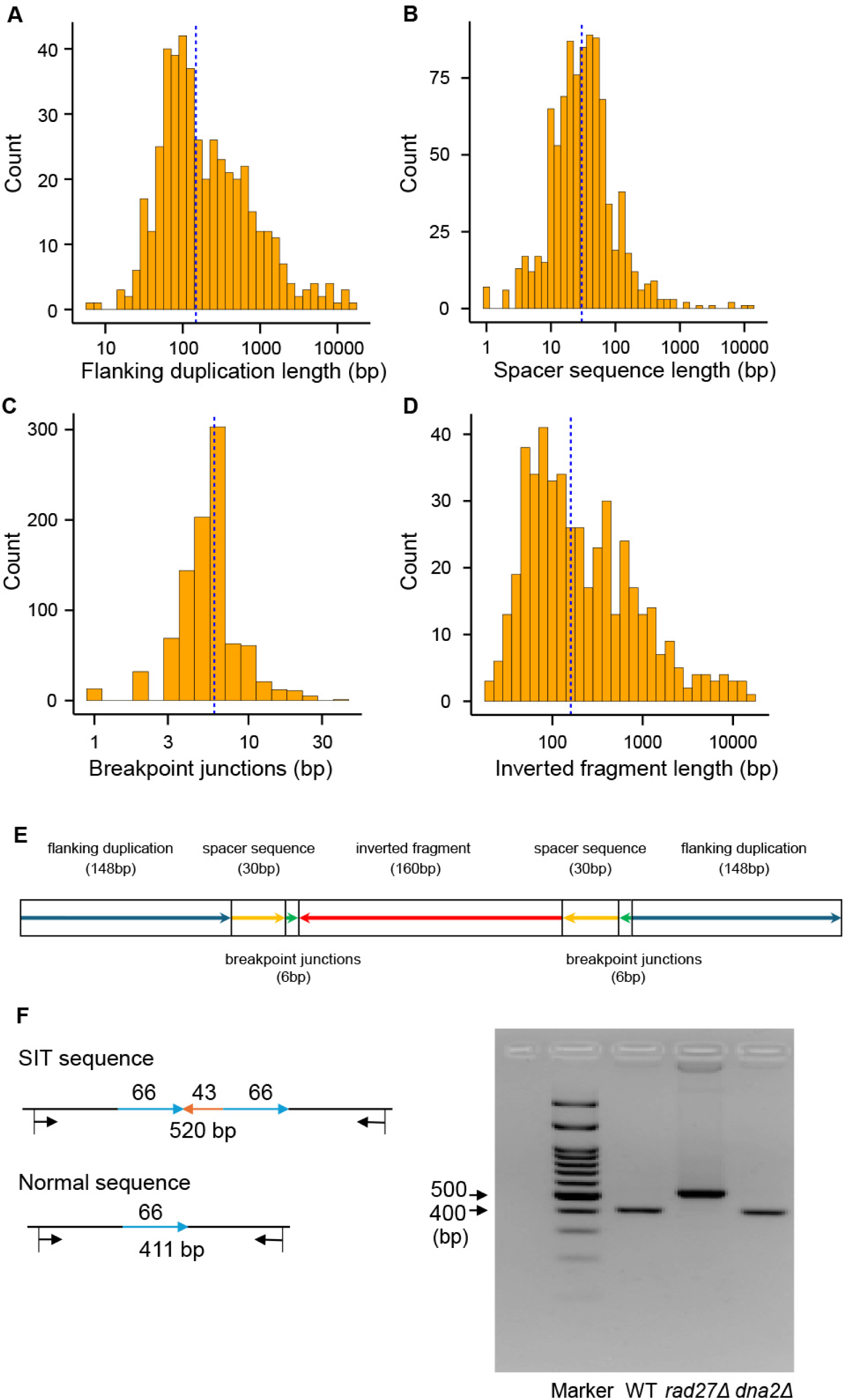
Structure features of SIT events identified in yeast strains. **(A)** Flanking duplications showed a broad size distribution ranging from 7 bp to > 10 Kbp, with a median of 148 bp. **(B)** Spacer fragments exhibited a narrow distribution, with most values clustering around a median of 30 bp. **(C)** Breakpoint junctions, characterized by reverse complementary sequences, displayed a sharply peaked distribution dominated by 6 bp. **(D)** Inverted fragments, forming the central reversed fragment, varied substantially in size from 20 bp to several Kbp, with a median length of 160 bp. **(E)** Schematic representation of a linear SIT structure, illustrating the flanking duplications, spacer sequence, breakpoint junctions, and inverted fragment. **(F)** PCR validation of a representative SIT event. The expected PCR products of the SIT sequence (520 bp) and the normal locus (411 bp) are shown.

Other genes in addition to *rad27* were found to be associated with high rates of global chromosomal rearrangement (GCR) in yeast.^[4]^ To identify additional contributors to SIT generation, we tested seven genes that are associated with high rates of GCR, including *dun1, mre11, rad1, sae2, mus81, rad51,* and *sgs1* (**Table 1** and **Table S8**). The results showed that one SIT event was detected in *dun1Δ, rad1Δ, sae2Δ,* and *sgs1Δ*; two events were detected in *exo1Δ* and *mus81Δ*; five events in *dna2-1*; and zero events in *mre11Δ*, *rad51Δ,* and *sgs1Δ* (**Table 1** and **Table S8**). This result suggests that all other tested genes are minor contributors to the generation of SIT events. Furthermore, we also examined SIT events in 15 human genomes (**Table S9**), none of which carried *FEN1* mutations, and no SITs were detected.

### 2.4. Genomic distribution and sequence features of SITs

We analyzed the genomic distribution of 484 SITs and found that they were widespread across the yeast genome (**Figure 3A** and **Table S8**). In the *rad27Δ* strain, two independent sequencing experiments identified 190 SITs in different loci with a few overlaps. The highest numbers of events occurred on chromosome (Chr) IV (15 and 16 events, respectively) while the fewest on Chr III (1 and 0 events) (**Figure S3** and **Table S8**). The *rad27Δ* strain maintained at 37 ℃ showed the highest frequency on Chr II (26 events), with the lowest observed on Chr I (1 event) (**Figure S3** and **Table S8**). It is interesting to note that in independent sequencing experiments, we obtained different event distribution patterns. However, there were hot spots where the events occurred frequently. We detected 11 recurrent sites across at least two independent experiments, including 8 sites detected twice, 1 site detected three times, and 1 site detected five times (**Figure 3A** and **Table S8**). Finally, among the 484 annotated inverted triplications, 407 overlapped with gene bodies, affecting 465 genes in total, including 84 essential genes representing 7.45% of all essential genes (*n* = 1,127) in yeast (**Table S8**).^52–54^

We next characterized the sequence features of SITs. The results showed that a complete SIT structure typically consists of four components: the flanking duplication, the spacer sequence, the breakpoint junctions, and the inverted fragment (**Figure 3A-E**). The flanking duplication showed a broad size distribution ranging from 7 bp to over 10 Kbp, with a median of 148 bp (**Figure 3A** and **Table S8**). The spacer fragment exhibited a narrow distribution, with most values clustering around a median of 30 bp (**Figure 3B** and **Table S8**). The breakpoint junctions are reverse complementary sequences^29^ with a sharply peaked distribution, dominated by 6 bp (**Figure 3C** and **Table S8**). The inverted fragment, which forms the central reversed DNA fragment, varied substantially in size from 20 bp to several kilobases, with a median length of 160 bp (**Figure 3D** and **Table S8**). Based on these data, we have sketched the linear structure of the SIT with a median size of 184/160/184 bp (DUP/IN/DUP) in **Figure 4E** and provided an SIT case that can be validated by PCR amplification in **Figure 4F**.

**Figure 4.**
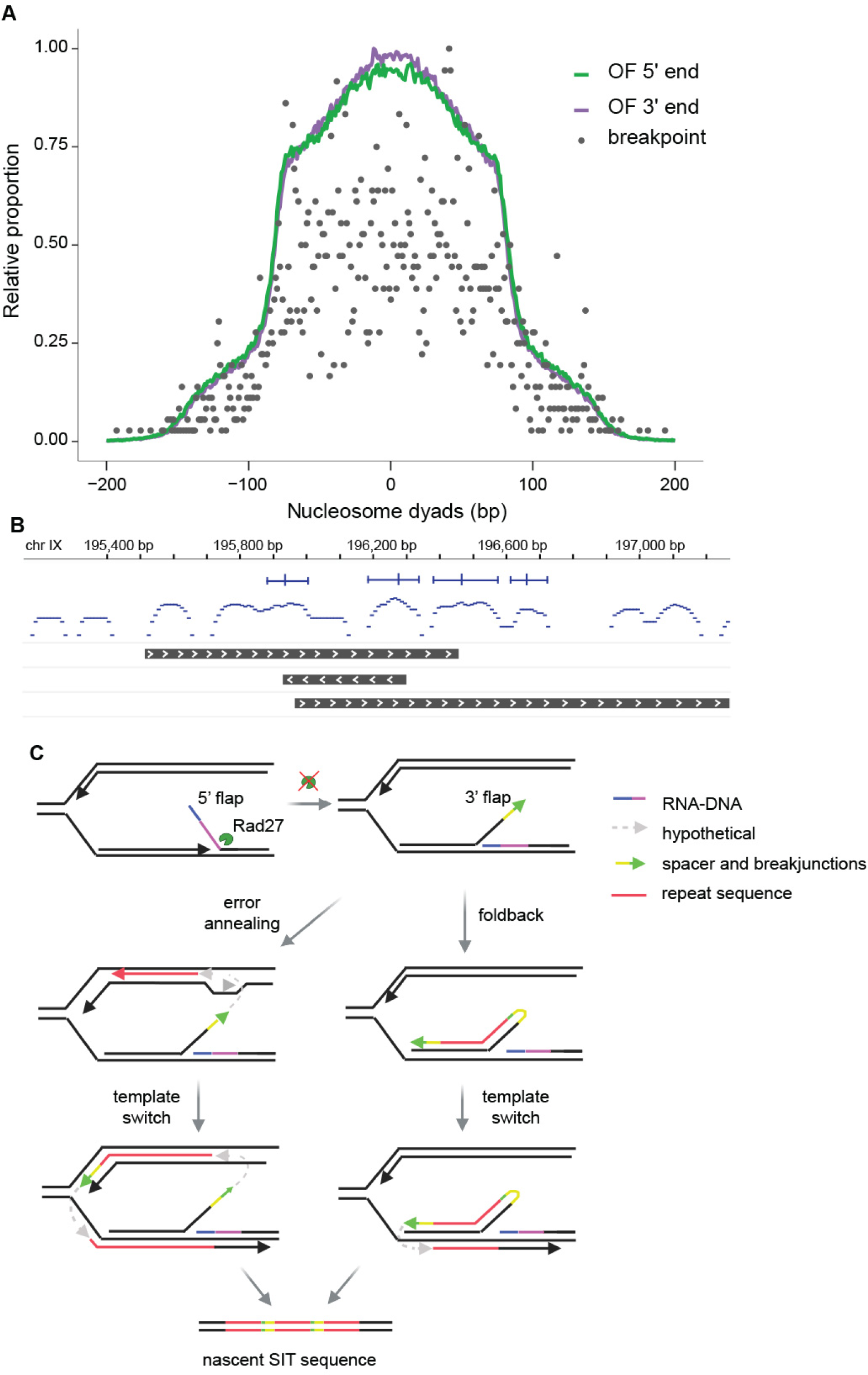
Breakpoint junctions of SITs are associated with nucleosomes and Okazaki fragment junctions. **(A)** Average distribution of Okazaki fragment 5’ ends, 3’ ends, and SIT breakpoint junctions relative to nucleosome dyads. **(B)** Genomic view of a representative SIT structure showing the breakpoint junctions mapped to nucleosome dyads. **(C)** Modeling SIT formation. When Rad27 is defective and 5’ flaps are not removed, 3’ flaps are formed. The 3’ flaps may either anneal to complementary sequences or fold back to prime DNA synthesis in the opposite direction, giving rise to an inverted fragment. Subsequent template switching at reverse complementary sequences promotes the nascent fragment to return to the original template and synthesize the DNA fragment one more time, producing a full SIT structure.

### 2.5. SIT breakpoint junctions are mapped to Okazaki fragment junction points

The association of major OFM enzyme FEN1 with SIT formation prompted us to investigate whether the SIT breakpoint junctions could be mapped to the Okazaki fragment termini. Previous studies have shown that Okazaki fragment termini preferentially occur near nucleosome midpoints (dyads) in yeast.^[55]^ Therefore, we performed micrococcal nuclease sequencing (MNase-seq) in both WT and *rad27Δ* strains, and identified 64,624 and 64,631 nucleosomes, respectively (**Table S10**, **Table S11**, and **Table S12**). Given that the stereotypical pattern of nucleosome depletion at promoters and well-ordered nucleosomes in gene bodies is found in all eukaryotes,^[56–60]^ we plotted the average nucleosome profile over all yeast genes (**Figure S4**). An overlay of hybridization profiles after alignment at the transcription start sites (TSS) showed an apparent nucleosome-depleted region (NFR) and a regular nucleosome array downstream of the TSS (**Figure S4**). Nucleosome signals around the transcript termination sites (TTS) were also defined, with a marked drop in coverage at the termination point followed by phased nucleosome occupancy downstream (**Figure S4**). Meanwhile, 19.81% of nucleosomes exhibited significant changes in position, occupancy, and fuzziness in *rad27Δ* compared to WT (FDR < 0.05; **Table S13**). We further integrated Okazaki fragment sequencing data,^[55]^ and the result showed that the breakpoint junctions of SITs, as well as Okazaki fragment termini, tend to show peak density around dyads (**Figure 4A-B** and **Table S14**).

Based on the data presented above, we propose that SITs are generated (**Figure 4C**) as follows: DNA polymerase δ displacement synthesis displaces the upstream 5’ flap for cleavage by Rad27. When Rad27 is defective, the 5’ flaps are not removed, and 3’ flaps arise, which normally invade the sister chromatin and prime DNA synthesis using the nascent DNA strand as a template. When the breakpoint junction sequence is palindromic, the 3’ flap may anneal to the complementary strand and prime DNA synthesis in the opposite direction, or the end sequence may fold back on itself, to form an inverted fragment. When the inverted synthesis meets resistance and finds another reverse complement sequence from the template strand of the original chromatin, the DNA polymerase synthesizes the original DNA fragment one more time.

### 2.6. Cellular instability and elimination of SIT

When the SIT structure was introduced in a plasmid to the *E. coli* system, the inverted fragment along with a copy of the duplicated fragment was progressively eliminated and recovered to the original locus sequence (**Figure 5A-B** and **Figure S5**). The SIT fragment density started to decrease by day 3, reaching ∼50% of the original density by day 4, with the ratio remaining relatively stable thereafter (**Figure 5B** and **Figure S5**). Given the rarity and possible genomic impact of SIT, we speculated that organisms may have evolved specific adaptive systems to cope with such rearrangements. We selected 24 genes with known functions in DNA replication, repair, and recombination as candidates to test this hypothesis (**Figure 5C**, **Figure S5**, and **Table S15**). With WT and 24 mutant *E. coli* strains, we show that loss of *holC* and *dnaQ* genes, which encode *E. coli* DNA polymerase III (Pol III) accessory subunits χ and ε, accelerated the SIT elimination processes significantly starting at day 0, even though the SIT band density rebounds beyond the WT density, which is probably due to induced expression of a compensatory gene for *dnaQ* (**Figure 5D** and **Figure S5**). Three other genes, including *rep,* which encodes the helicase Rep, and nuclease subunits *sbcC* and *sbcD*, which form the SbcCD structure-specific nuclease complex, tolerated SIT mutations until Day 15 without popping it out (**Figure 5C-D** and **Figure S5**).

**Figure 5.**
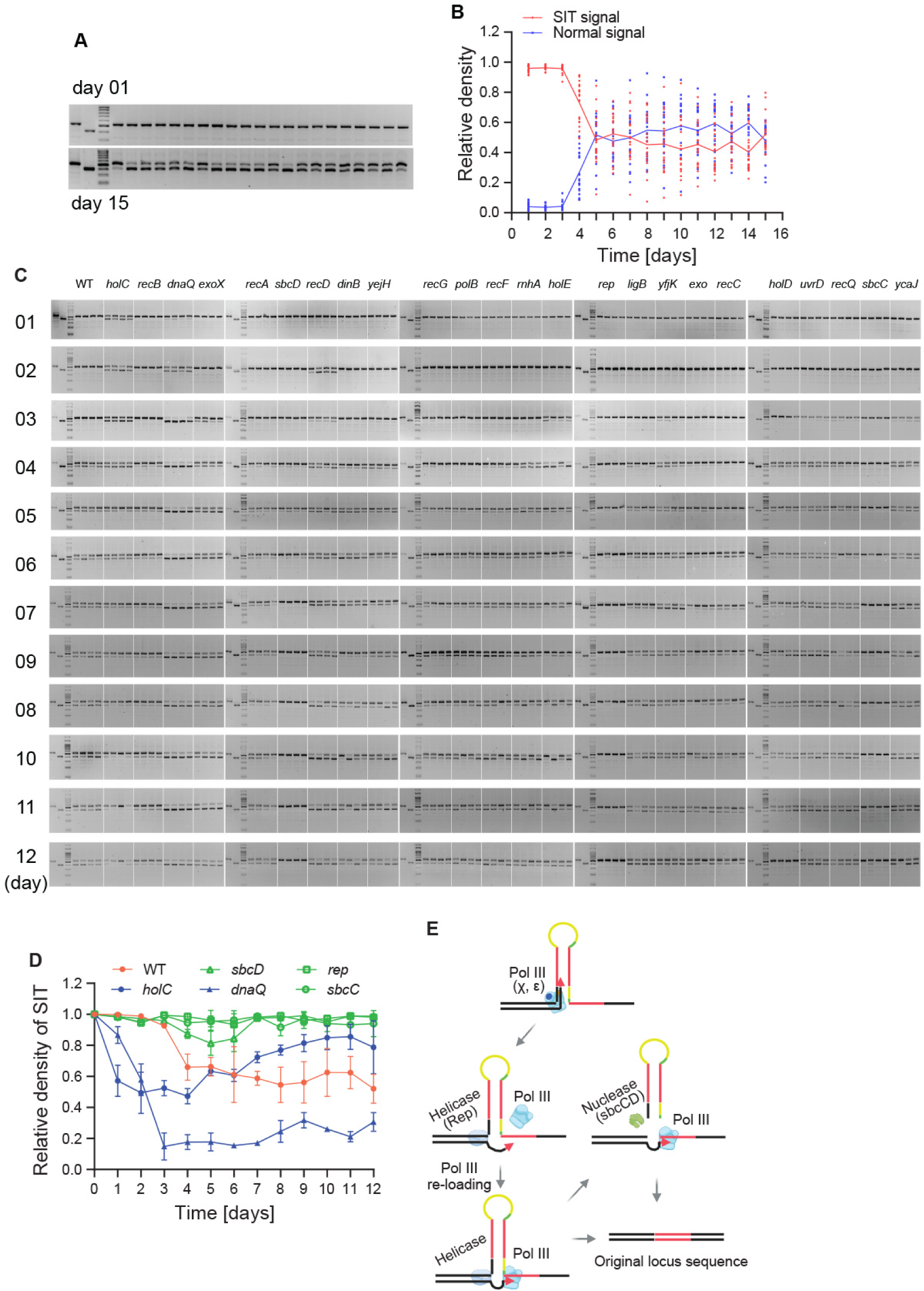
Dynamic equilibrium of SIT structure in cells. **(A)** and **(B)**, Quantification of signal dynamics of the normal (blue) and SIT (red) in the WT strain over time. **(C)** PCR-based monitoring of SIT and normal structure signals over 12 days following plasmid transformation in WT and 24 mutants (**Figure S5**). **(D)** Relative density of SIT signal in WT and five representative mutants over time, showing their effects on the SIT structure stability (**Figure S5**). Error bar, mean ± standard deviation. **(E)** Proposed mechanism depicting how the SIT structures are precisely eliminated by DNA Pol III slippage, helicase displacement of the primer, and structure-specific nuclease cleavage of hairpin structures.

Based on this set of data, we propose a model (**Figure 5E**) on how the SIT structure is eliminated: *E. coli* DNA Pol III accessory subunits χ and ε enable displacement synthesis of the hairpin structure and replicate the full length of the SIT mutation. Knock out of these two factors leads to hairpin (made of DUP-IN, spacer, and breakpoint junctions) slippage and accelerated elimination of hairpin structures. Rep helicase separates the synthesized DNA from the hairpin and promotes the slippage process. Meanwhile, the knockout of *sbcC* or *sbcD* inactivates the hairpin-cleaving activity of the SbcCD endonuclease complex,^[61]^ which leads to stabilization of the SIT structure.

## 3. Discussion

SVs are one of the major classes of genetic variation and are closely associated with evolution^[62–64]^ and human disease.^[65]^ The discovery and characterization of SVs, interpreting their genetic consequences, have been a central focus of genetic research.^[5]^ In this study, we describe a new type of SV characterized by a DUP/IN/DUP structure, a spacer sequence, and breakpoint junctions, with median lengths of 184/160/184, 30, and 6 bp, respectively. Although they share structural features with locus-specific inverted triplications, which vary from kilobase to megabase in previously reported cases,^[40,41]^ SITs are clearly distinguished by their substantially smaller size and unique mechanism of formation.

The observed low frequency of SIT events in human cancer genomes, with 745 samples showing no detectable SITs, may reflect both the inherent rarity and the profound impact that such complex SVs can exert on genome architecture and gene function. While short-read sequencing technology lacks the resolution to capture many complex SVs, particularly in repetitive or complex genomic regions,^[66–70]^ long-read PacBio HiFi sequencing offers clear advantages for complex SV detection,^[68,71]^ and our PacBioR software, specifically optimized for this platform, outperforms existing tools in identifying complex SVs, particularly SITs. Thus, integrating PacBio HiFi sequencing with PacBioR may help address these limitations and facilitate genome-wide analyses of rare and complex SVs.

Among DNA-associated genes, FEN1 mutations exhibited the highest OR enrichment, highlighting a close association between FEN1 and SIT formation. This was further validated in yeast, where deletion of the FEN1 yeast homologue *rad27Δ* resulted in a substantial increase in SIT events. The ODIRA model, based on the observation that the inverted triplication contains the origin of replication (*ARS228*), proposes that replication errors at origins, followed by fork reversal and ligation, drive the formation of inverted triplications.^[27,41,42]^ In addition, Carvalho et al. proposed that the inverted triplication can arise from the restart of a collapsed replication fork, in which a DNA break generates a 3′ tail that invades the opposite sister strand.^[29,40]^ Therefore, we established a predicted model attributing the origination of SIT structure to DNA replication and functional deficiency of the major OFM enzyme, FEN1/Rad27. The unprocessed 5’ flap blocks the DNA polymerase δ from displacement synthesis and leads to the formation of a 3’ flap, which is capable of sister chromatin strand invasions.^[50,51]^ When the 3’ flap possesses a palindromic sequence motif, it may invade and anneal with the complementary sequence in the opposite direction. The newly synthesized DNA in the opposite direction must undergo a second template switch, triggered by fork stalling on the original template strand, which has been documented previously.^[27,29,40–42]^ Alternatively, the 3’ flap may fold back on itself when the complementary sequence is available^[50,51]^ and have a template switch at the upstream nick to return DNA synthesis to the original template, generating a SIT structure.

Our study demonstrated the inherent instability and dynamic equilibrium of SIT structures in a cellular context. The functional deficiency of SbcCD suppresses the resolution of inverted triplications, indicating that these rearrangements may form hairpin structures in the cell that are specifically recognized and cleaved by hairpin-resolving nucleases.^[61]^ Meanwhile, the presence of both an inverted repeat and direct repeat (DR) within an inverted triplication may provide a potential substrate for replication slippage.^[72–74]^ When replicating a DR, the polymerase may stall and dissociate from the newly synthesized strand, then realign with the second duplication and resume synthesis, causing additions or deletions of repeat units.^[72,74]^ *In vitro* studies have shown that the loss of *E. coli* 3’ → 5’ exonuclease, polymerase II, and the T4 and Φ29 phage polymerases, is correlated with the inhibition of replication slippage.^[74]^ However, our study showed that functional deficiency of DnaQ, which primarily contributes to the 3′→5′ exonuclease activity of DNA Pol III, promoted the resolution of inverted triplications relative to WT. This discrepancy may be attributed to differences in experimental systems or enzymatic contexts. Our results indicated that the deletion of *holC*, which stabilizes DNA polymerase III on single-stranded DNA via interaction with single-stranded DNA-binding protein, accelerated the repair of inverted triplications. This finding is consistent with previous studies showing that single-stranded DNA-binding proteins can suppress the occurrence of replication slippage.^[74]^ In addition, we found that slippage is likely promoted by the helicase activity of Rep. When the polymerase stalls, Rep may bind to the end of the nascent strand end and initiate unwinding along the DNA in the 3′ to 5′ direction,^[75–77]^ thereby facilitating strand dissociation during the slippage process.

In summary, the SIT structure as a novel SV has been characterized in this study. The PacBioR package offers a robust and scalable method for detecting complex SVs, particularly SIT structures. Our mechanistic analyses demonstrate that replication errors during OFM are a major driver of SIT formation and provide two possible pathways underlying its origin. We further propose two repair routes for the SIT structure, replication slippage and hairpin cleavage, which together help explain the inherent instability and dynamic maintenance of SIT structures *in vivo*. The rarity of SITs likely reflects both the stringent conditions required for their formation and evolved cellular repair systems, whereas imperfect repair may account for the frequent co-occurrence of other SVs at the same loci.^[29,40,42]^ This study also has limitations, as our mechanistic dissection was confined to yeast and *E. coli*, and its applicability to mammalian systems remains to be established. In addition, the molecular events driving the second template switch during SIT formation remain unresolved. Addressing these gaps is essential for the refinement of our model and further clarifying the role of SITs in genome stability and tumorigenesis.

## 4. Experimental Methods

### Cancer Genome Datasets Collection and Analysis

We collected 1,340 short-read sequencing datasets, including both 739 whole-genome sequencing (WGS) and 591 whole-exome sequencing (WES) samples, derived from 29 projects covering 22 distinct cancer types (**Figure 1A** and **Table S1**), for use in this study. All short-read sequencing datasets used in this study were obtained from the NCBI Sequence Read Archive (https://www.ncbi.nlm.nih.gov/sra/), except for the data from B-ALL patients, which were generated by our laboratory (**Table S1**). Paired-end reads containing adapters, ploy-N, or any bases with Phred quality scores lower than 20 were removed with Cutadapt v.2.1 and TrimGalore v.0.6.5 (https://www.bioinformatics.babraham.ac.uk/projects/trim_galore/). The clean reads were aligned to the hg38 reference genome using Bowtie2 v.2.2.9.^[78]^ Duplicated reads were removed by the Picard v.2.23.4 MarkDuplicates module (https://broadinstitute.github.io/picard/index.html). The Pindel v.0.2.5^[12]^ was used to call SVs from the alignment data. We filtered the Pindel results by requiring the length of the alternative sequence (ALT) to be more than twice that of the reference sequence (REF) in one SV. Then the alternative sequences were re-aligned to the reference genome using Blastn v.2.16.0,^[79]^ and candidate SITs were filtered by identifying alignment patterns featuring a central reverse-oriented segment flanked by two direct fragments. The SITs were subsequently confirmed through manual inspection using Integrative Genomics Viewer (IGV)^[80]^ (**Figure S1A**). The SNVs and indels were identified by VarScan2 v.2.3.7.^[81]^ All variants were annotated using Annovar (http://www.openbioinformatics.org/annovar/).^[82]^

### Yeast Strains and PacBio HiFi Sequencing

The WT yeast used in the study was the haploid MATa RDKY2669 strain (**Table S7**). Single gene deletion derivatives of WT, including *rad27*, *rad1*, *rad51*, *dun1*, *mre11*, *mus81*, *sgs1*, *sae2*, were generated via a homologous recombination-based strategy. WT cells were first transformed with PCR-amplified DNA fragments containing *his3*, *trp1*, or *ura3* as selection markers, flanked by 39 bp homologous arms corresponding to the target loci (**Table S7** and **Table S16**). Stable transformants were then selected using corresponding auxotrophic plates, and successful gene knockout was confirmed by Sanger DNA sequencing. The *dna2-1* and *exo1* mutant strains were gifted by Judith L. Campbell (California Institute of Technology).

Unless otherwise specified, all yeast strains were cultured in YPD medium at 30 °C with shaking at 200 rpm. Yeast cells were inoculated at an initial OD600 of 0.1 into liquid YPD medium and grown overnight, and then DNA was extracted using the YeaStar™ Genomic DNA Kit (ZYMO Research, D2002) according to the manufacturer’s instructions. After confirming the quality of the genomic DNA, the SMRTbell libraries were prepared with SMRTbell Prep Kit 3.0 (Pacific Biosciences) following the manufacturer’s protocol. Briefly, The Megarupter 3 system (Diagenode, Denville, NJ, USA) was used for genomic DNA shearing, and the Blue Pippin system (Sage Science, Beverly, MA, USA) was used for size-selection to keep fragments greater than 13 Kbp. A subset of the libraries was constructed using barcoded adapters to allow pooling before size-selection. Each library was run on one SMRT Cell (version 8M) using version 2.0 sequencing reagents. The Sequel IIe sequencer (Pacific Biosciences, Menlo Park, CA, USA) instrument was used to generate the raw subreads. The circular consensus sequences were extracted from the subreads and then filtered for short (< 100 bp) or low quality (QV < 80) using the SMRTLink v. 9.0.0.92188 package to produce the HiFi reads. Samples treated at 37 °C were initially cultured at 30 °C, then incubated at 37 °C for 4 hours, with all subsequent procedures conducted in parallel with untreated samples.

### Development of PacBioR for Detecting SVs Using HiFi Reads

To improve the sensitivity of inverted triplication detection, we have implemented an algorithm called “PacBioR” using an accurate alignment and rule-based SV calling scheme (**Figure 2**). The reads were first aligned to a reference genome using pbmm2 (https://github.com/PacificBiosciences/pbmm2). The alignments were then examined for long stretches of insertion, deletion, or soft clips. Reads with these abnormalities were considered candidates for further analysis. The MegaBLAST algorithm^[83]^ was used to realign the candidates with improved accuracy. To overcome its slow speed, we utilized a high-speed version, hs-blastn,^[84]^ which is more than 20× faster than regular megaBLAST. Several optimization steps were implemented to resolve the complex alignment structure. These steps included: resolving multiple alignment regions among distal regions, removing redundant alignments within a local region, and filtering out the alignments with a large number of mismatches. These cleaned alignment results were then converted into two vectors, one representing the alignment along the query read and the other representing the alignment along the reference genome. This can be achieved by projecting the aligned segments onto query and reference sequences, with 1 representing a match, 0 representing a missing base, and −1 representing a match on the opposite strand. The two vectors serve as the foundation for identifying candidate SVs. For example, if the query vector contains a stretch of 0 and the reference vector contains no 0, the read contains an insertion. Deletion events will have the opposite pattern. Duplications are represented by a stretch of integers greater than or equal to 2 in the query vector, while the reference vector contains only 1. Inversions have a stretch of −1 in the query. Inverted duplications can be indicated by both 1 and −1 stretches in the query and a stretch of integers larger than 2 in the reference. This detection process was repeated in a sliding window approach to cover all segments in each query read, ensuring multiple SV events on each read could be detected as well. Each read was examined independently and assigned an SV event based on the processes outlined above. Similar SV events were then aggregated, and the number of supporting reads was recorded along with the total coverage at these loci.

### MNase-Seq and Data Analysis

The mono-nucleosomes were extracted from WT or *rad27Δ* yeast cells (**Table S9**) according to a previously described protocol^[85]^ with a small modification. Cells were grown in YPD medium in stable log phase (OD600 ≈ 0.6) and harvested by centrifugation. The cells were fixed with formaldehyde at a final concentration of 1% for 15 min, followed by quenching with glycine at a final concentration of 125 mM for 5 min. The cells were then pelleted and washed twice with cold sterile water. The pellet was resuspended with lysis buffer (1 M sorbitol, 50 mM Tris–HCl, pH 7.4, 10 mM *ß*-mercaptoethanol), and zymolyase solution (10 mg/mL zymolyase dissolved in 40% glycerol, 60% lysis buffer) was added at a final concentration of 0.25 mg/mL. Cells were incubated at 30 °C with gentle shaking, and then the spheroplasts were collected by centrifugation at 3000 *g* for 10 min at 4 °C. The spheroplasts were gently resuspended in resuspension buffer (0.5 mM spermidine, 0.075% NP-40, 50 mM NaCl, 10 mM Tris–Cl, pH 7.5, 5 mM MgCl₂, 1 mM CaCl₂, 1 mM β-mercaptoethanol) and digested with micrococcal nuclease (300 U/μL) at 37 °C for 30 min. The reaction was stopped on ice by adding EDTA to a final concentration of 500 mM. DNA was purified using phenol–chloroform extraction and concentrated by ethanol precipitation. The integrity and size distribution of the digested fragments were determined using the microfluidics-based platform Bioanalyzer (Agilent). A PCR-free protocol with the KAPA Library Preparation kit (Roche) was employed to prepare short-insert paired-end libraries for MNase sequencing. The libraries were sequenced using TruSeq SBS Kit v4-HS (Illumina) following the manufacturer’s protocol. The raw reads were quality checked by FastQC v.0.20.0 (https://www.bioinformatics.babraham.ac.uk/projects/fastqc/). The reads containing adapters, ploy-N, or any bases with Phred quality scores lower than 20 were removed with Cutadapt v.2.1 and TrimGalore v.0.6.5 to generate clean reads. The reads were then aligned to the reference yeast genome sacCer3 using Bowtie2 v2.2.9. Nucleosome positioning and occupancy were examined by DANPOS2 v.2.2.2^[86]^ using the function Dpos.

### Human HiFi Sequencing Data Collection

Leukemic cells from the bone marrow aspirates of three patients diagnosed with B-ALL were collected at diagnosis and isolated by density gradient centrifugation. Samples of < 70% tumor cell content were purified by fluorescence-activated cell sorting. Genomic DNA was extracted from the cells using a commercial DNA extraction kit (PureLink Genomic DNA Mini Kit, Invitrogen, Carlsbad, CA, USA). After quality check of the genomic DNA, HiFi library preparation and sequencing were performed following the same protocol used for yeast samples. In addition, we incorporated 12 publicly available HiFi sequencing datasets into our study, including data derived from breast carcinoma, melanoma, peripheral blood, lymphoblastoid cells, and the Human Genome Structural Variation Consortium (**Table S9**).

### *E. coli*-based Functional Screening of Factors Involved in Inverted Triplication Stability

PCR amplification of normal and inverted triplication fragments was performed using genomic DNA, 12.5 μL of 2× Taq PCR Premix (Bioland Scientific), and 0.2 µM of forward and reverse primers (**Table S15**). PCR conditions were 95 °C for 5 min followed by 35 cycles of 95 °C, 5 sec; 58 °C, 30 sec; 72 °C, 45 sec; with a final elongation step of 72 °C for 10 min. The PCR-amplified inverted fragments were ligated into TOPO TA® cloning vectors and subsequently transformed into *E. coli* wild-type and 23 single-gene knockout strains (Horizon, catalog#OEC4987) (**Table S14**). Transformed cells were plated on LB agar supplemented with ampicillin (100 μg/mL) and grown overnight at 37 °C. Single colonies were picked and cultured into 4 mL of LB liquid medium supplemented with ampicillin (37 ℃, 270 rpm). After 24 hours, 4 μL of the culture was transferred into fresh 4 mL LB medium under the same conditions. After 24 hours of growth, 4μL of the bacterial culture was transferred into 4 mL of fresh medium. This process was repeated under the same conditions for sequential subcultures. At each passage, PCR analysis was conducted to evaluate the status of both the normal and inverted triplication fragments.

## Acknowledgements

Acknowledgements

We gratefully acknowledge the Sequencing Core and the Computational Data Center at City of Hope for their technical support. We additionally thank Judith L. Campbell (California Institute of Technology) and Richard Kolodner (University of California, San Diego) for providing the yeast strains and the members of Zhaohui Gu’s laboratory for their assistance in acquiring human samples.

## Data Availability Statement

Sequence data have been deposited at NCBI as Submission ID PRJNA1348743 and are publicly available as of the date of publication. Original code for PacBioR has been deposited at GitHub and is publicly available at https://github.com/xiweiwu/pacbior as of the date of publication. All other data reported in this paper will be shared by the corresponding authors upon request.

---

**Figure S1.**
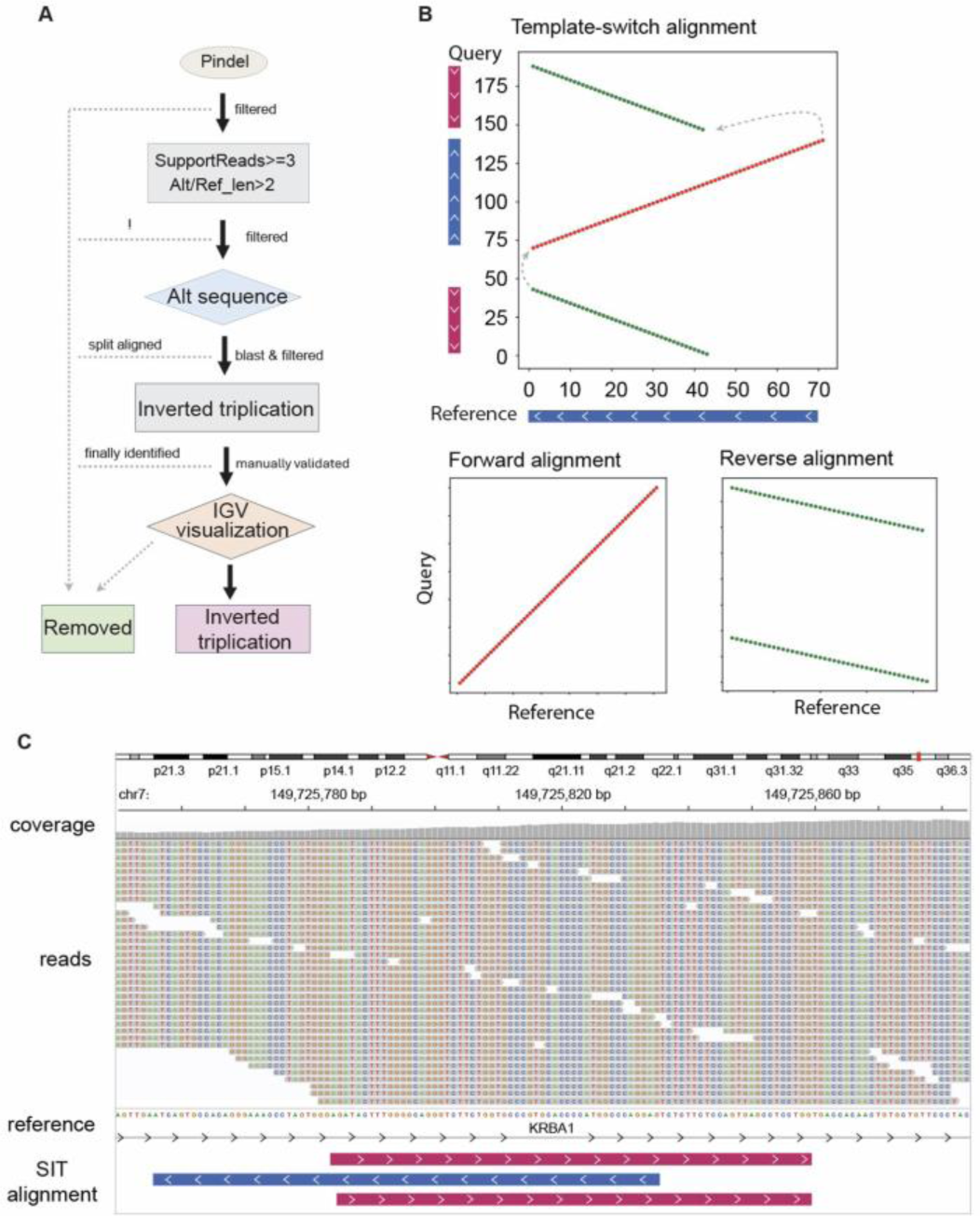
Workflow for the detection and validation of SIT events using short-read sequencing data. **(A)** Pipeline for SIT event identification. SVs were called with Pindel and filtered to retain only SVs with ≥ 3 supporting reads and the alternative/reference length ratio > 2. The candidate SVs were realigned using Blastn to generate the predicted SIT events. These SITs were validated through IGV visualization to confirm the characteristic structure, defined by a central inverted segment flanked by two direct copies. **(B)** Template-switch alignment of SIT structure. The top panel shows a template-switch alignment plot, where red segments represent forward alignment and green segments represent reverse alignments. The bottom panels display separate alignments: the forward alignment (left, red) corresponding to the inverted segment, and the reverse alignment (right, green) corresponding to the direct copies. **(C)** IGV visualization of a SIT event resolved by split alignment to the reference genome.

**Figure S2.**
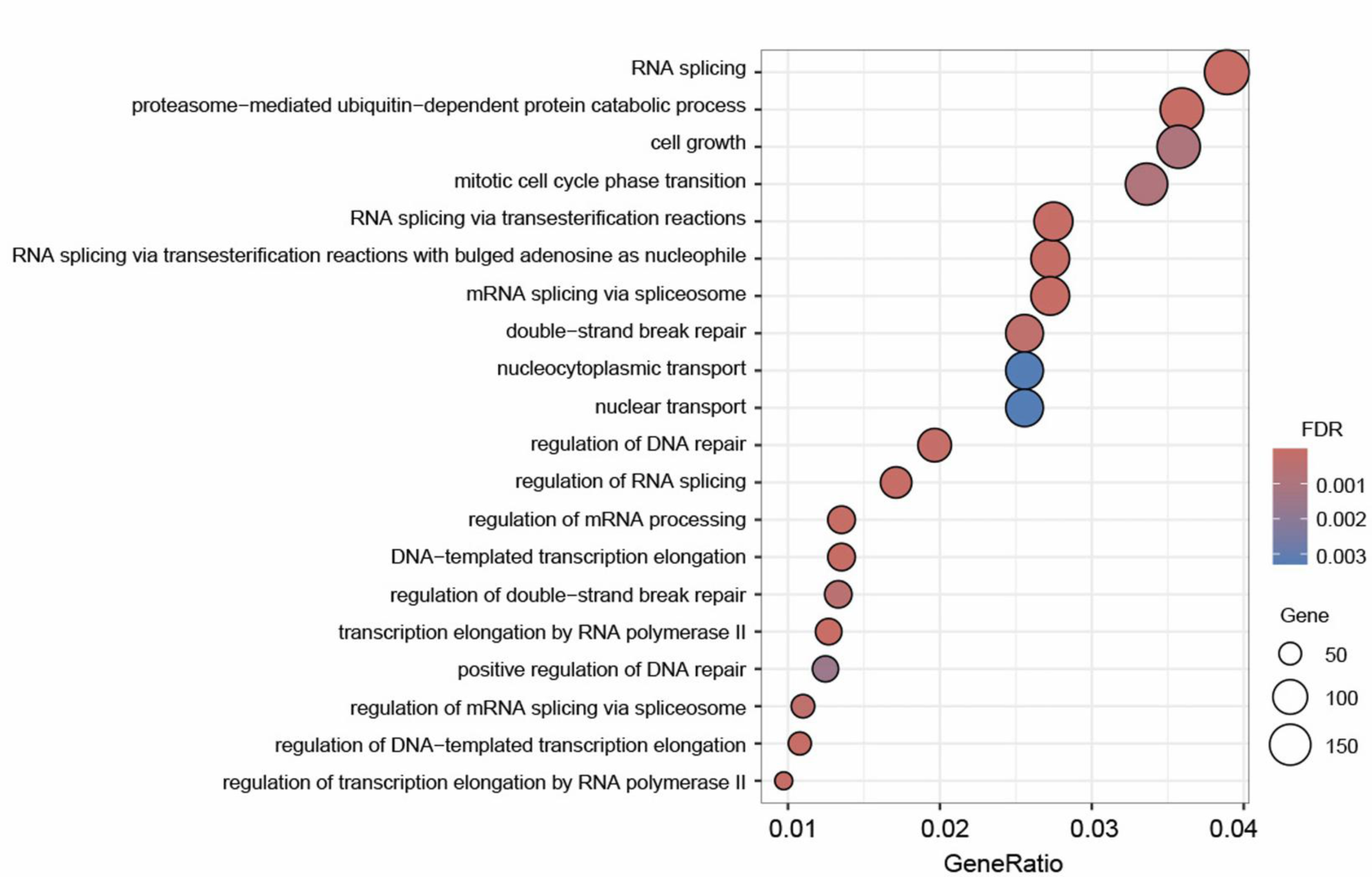
The top 20 significant GO biological processes enriched in genes that were significantly enriched in the high SIT incidence group. Go analysis showed that these enriched genes are involved in nucleic acid metabolism, including RNA splicing, DNA replication, recombination, and repair (Table S4).

**Figure S3.**
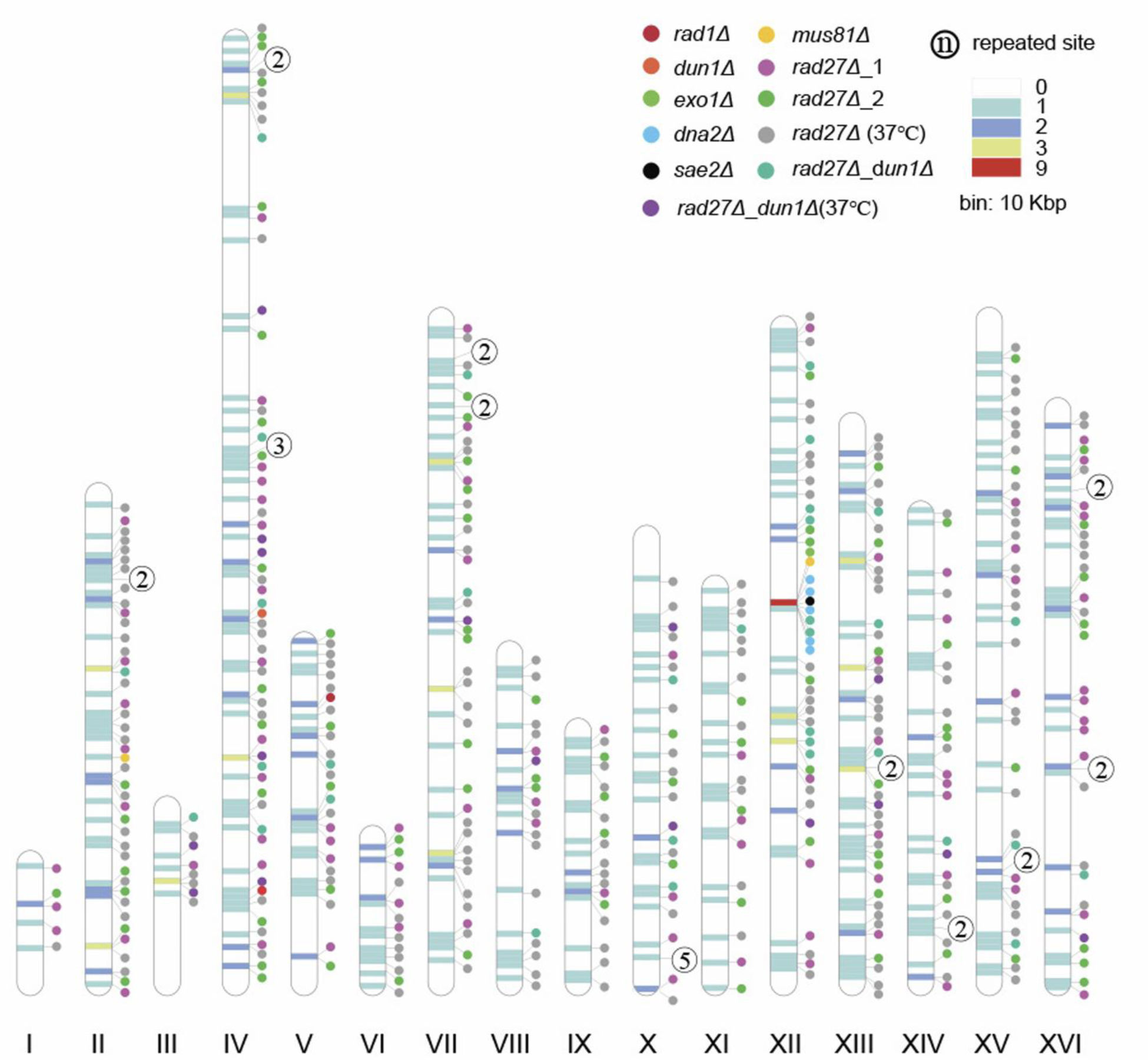
Genome-wide distribution of SIT events in yeast strains. Each dot represents a SIT event, color-coded by yeast strain and experimental conditions. “repeated site” indicates how many of the 17 sequencing datasets detected the SIT event at the same locus.

**Figure S4.**
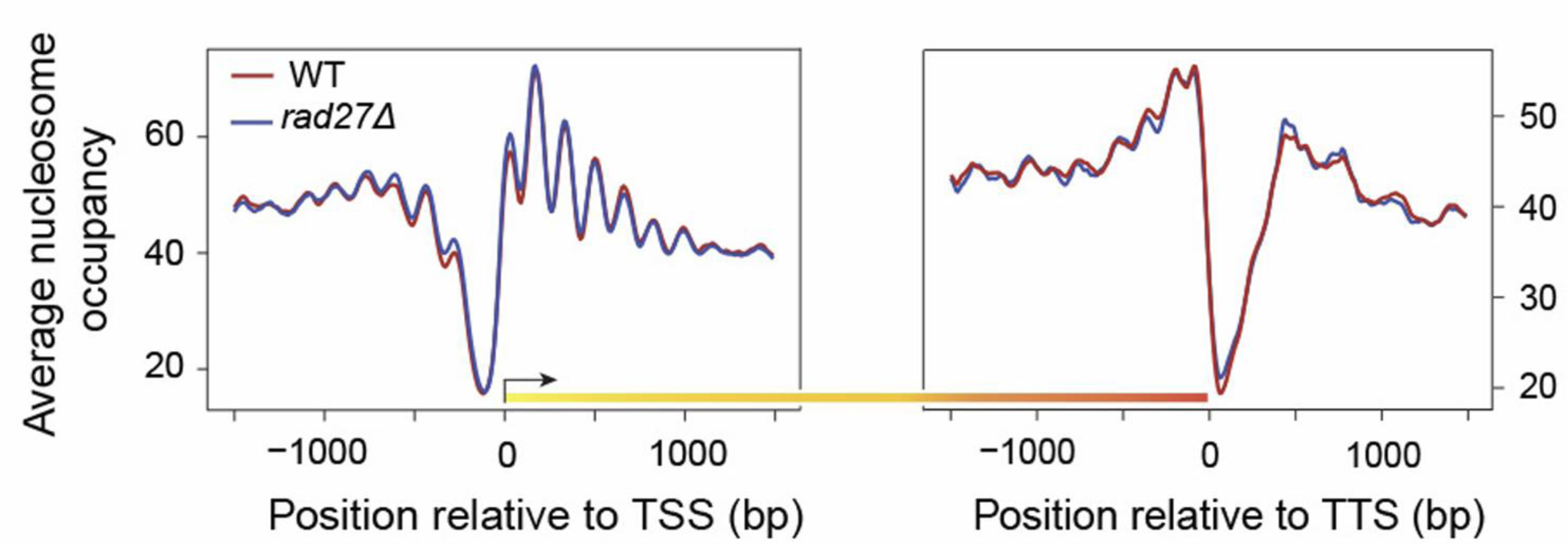
Nucleosome occupancy profiles aligned at transcription start sites (TSS) and transcription termination sites (TTS). The left panel shows a nucleosome-depleted region (NDR) at the TSS, followed by a regular phased nucleosome array downstream. The right panel shows nucleosome occupancy around the TTS, with a sharp drop in coverage at the termination point and phased nucleosome positioning downstream.

**Figure S5.**
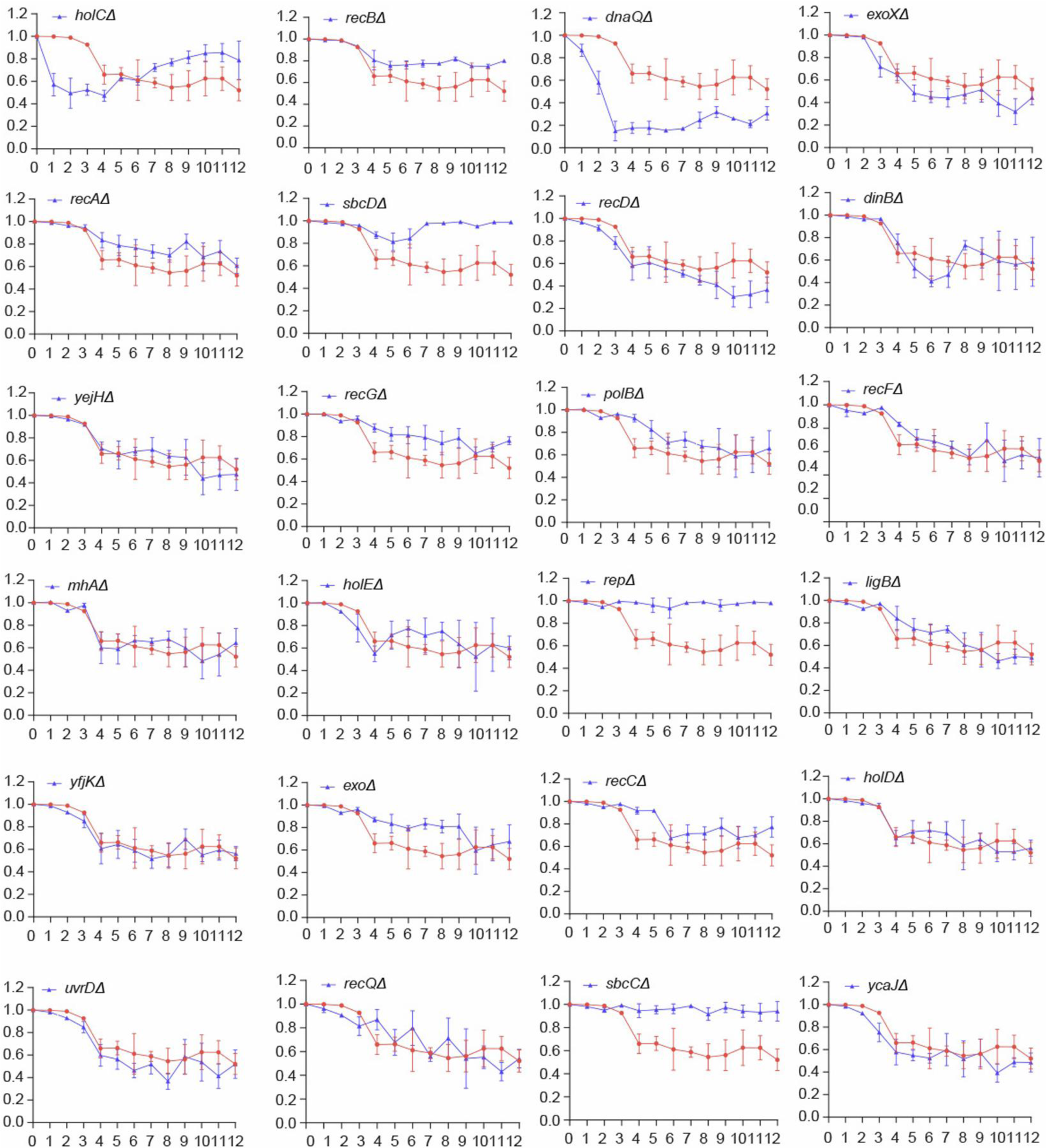
Quantification of SIT structure repair in 24 E. coli mutants compared to WT. Each panel shows the time course quantification of SIT intensity in a mutant (red) compared with WT (blue) over a 12-day period. Error bar, mean ± standard deviation.

## References

[1] Stankiewicz, P. & Lupski, J. R. (2010). Structural variation in the human genome and its role in disease. Annu. Rev. Med. 61, 437–455.

[2] Feuk, L., Carson, A. R. & Scherer, S. W. (2006). Structural variation in the human genome. Nat. Rev. Genet. 7, 85–97.

[3] Hollox, E. J., Zuccherato, L. W. & Tucci, S. (2022). Genome structural variation in human evolution. Trends Genet. 38, 45–58.

[4] Putnam, C. D. & Kolodner, R. D. (2017). Pathways and mechanisms that prevent genome instability in *Saccharomyces cerevisiae*. Genetics 206, 1187–1225.

[5] Collins, R. L. & Talkowski, M. E. (2025). Diversity and consequences of structural variation in the human genome. Nat. Rev. Genet. 26, 443–462.

[6] Trask, B. J. (2002). Human cytogenetics: 46 chromosomes, 46 years and counting. Nat. Rev. Genet. 3, 769–778.

[7] Heng, H. H., Squire, J. & Tsui, L. C. (1992). High-resolution mapping of mammalian genes by in situ hybridization to free chromatin. Proc. Natl. Acad. Sci. 89, 9509–9513.

[8] Chester, M. et al. (2012). Extensive chromosomal variation in a recently formed natural allopolyploid species, *tragopogon miscellus* (asteraceae). Proc. Natl. Acad. Sci. 109, 1176–1181.

[9] Vissers, L. E. L. M. et al. (2003). Array-based comparative genomic hybridization for the genomewide detection of submicroscopic chromosomal abnormalities. Am. J. Hum. Genet. 73, 1261–1270.

[10] Yoon, S., Xuan, Z., Makarov, V., Ye, K. & Sebat, J. (2009). Sensitive and accurate detection of copy number variants using read depth of coverage. Genome Res. 19, 1586–1592.

[11] Chen, K. et al. (2009). BreakDancer: an algorithm for high-resolution mapping of genomic structural variation. Nat. Methods 6, 677–681.

[12] Ye, K., Schulz, M. H., Long, Q., Apweiler, R. & Ning, Z. (2009). Pindel: a pattern growth approach to detect break points of large deletions and medium sized insertions from paired-end short reads. Bioinformatics 25, 2865–2871.

[13] Chen, K. et al. (2014). TIGRA: A targeted iterative graph routing assembler for breakpoint assembly. Genome Res. 24, 310–317.

[14] Hormozdiari, F. et al. (2010). Next-generation VariationHunter: combinatorial algorithms for transposon insertion discovery. Bioinformatics 26, i350–i357.

[15] Jiang, Y., Wang, Y. & Brudno, M. (2012). PRISM: pair-read informed split-read mapping for base-pair level detection of insertion, deletion and structural variants. Bioinformatics 28, 2576–2583.

[16] Rausch, T. et al. (2012). DELLY: structural variant discovery by integrated paired-end and split-read analysis. Bioinformatics 28, i333–i339.

[17] Sudmant, P. H. et al. (2015). An integrated map of structural variation in 2,504 human genomes. Nature 526, 75–81.

[18] Durbin, R. M. et al. (2010). A map of human genome variation from population-scale sequencing. Nature 467, 1061–1073.

[19] Pollard, M. O., Gurdasani, D., Mentzer, A. J., Porter, T. & Sandhu, M. S. (2018). Long reads: their purpose and place. Hum. Mol. Genet. 27, R234–R241.

[20] Sedlazeck, F. J., Lee, H., Darby, C. A. & Schatz, M. C. (2018). Piercing the dark matter: bioinformatics of long-range sequencing and mapping. Nat. Rev. Genet. 19, 329–346.

[21] Burgess, D. J. (2018). Genomics: Next regeneration sequencing for reference genomes. Nat. Rev. Genet. 19, 125–126.

[22] Jiang, T. et al. (2020). Long-read-based human genomic structural variation detection with cuteSV. Genome Biol. 21, 189.

[23] Qi, W. et al. (2022). The haplotype-resolved chromosome pairs of a heterozygous diploid African cassava cultivar reveal novel pan-genome and allele-specific transcriptome features. Gigascience 11, giac028.

[24] Lin, J. et al. (2022). SVision: a deep learning approach to resolve complex structural variants. Nat. Methods 19, 1230–1233.

[25] Liu, E. et al. (2023). Unraveling Diverse Mechanisms of Complex Structural Variant Interactions through Multiomic Data in Multiple Myeloma. Blood 142, 641.

[26] Mahmoud, M. et al. (2019). Structural variant calling: the long and the short of it. Genome Biol. 20, 246.

[27] Brewer, B. J. et al. (2015). Origin-dependent inverted-repeat amplification: tests of a model for inverted DNA amplification. PLOS Genet. 11, e1005699.

[28] C., Martin, S., Trask, B. & Hamlin, J. L. (1993). Sister chromatid fusion initiates amplification of the dihydrofolate reductase gene in Chinese hamster cells. Genes Dev. 7, 605–620.

[29] Grochowski, C. M. et al. (2024). Inverted triplications formed by iterative template switches generate structural variant diversity at genomic disorder loci. Cell Genomics 4, 100590.

[30] Giorda, R. et al. (2007). Two classes of low-copy repeats comediate a new recurrent rearrangement consisting of duplication at 8p23.1 and triplication at 8p23.2. Hum. Mutat. 28, 459–468.

[31] Soler-Alfonso, C. et al. (2014). CHRNA7 triplication associated with cognitive impairment and neuropsychiatric phenotypes in a three-generation pedigree. Eur. J. Hum. Genet. 22, 1071–1076.

[32] Beri, S., Bonaglia, M. C. & Giorda, R. (2013). Low-copy repeats at the human VIPR2 gene predispose to recurrent and nonrecurrent rearrangements. Eur. J. Hum. Genet. 21, 757–761.

[33] Zafar, F. et al. (2018). Genetic fine-mapping of the Iowan SNCA gene triplication in a patient with Parkinson’s disease. NPJ Parkinsons Dis. 4, 1–7.

[34] Devriendt, K., Matthijs, G., Holvoet, M., Schoenmakers, E. & Fryns, J.-P. (1999). Triplication of distal chromosome 10q. J. Med. Genet. 36, 242–245.

[35] Schinzel, A. A. et al. (1994). Intrachromosomal triplication of 15q11-q13. J. Med. Genet. 31, 798–803.

[36] Rivera, H., Bobadilla, L., Rolon, A., Kunz, J. & Crolla, J. A. (1998). Intrachromosomal triplication of distal 7p. J. Med. Genet. 35, 78–80.

[37] Eckel, H. et al. (2006). Intrachromosomal triplication 12p11.22–p12.3 and gonadal mosaicism of partial tetrasomy 12p,. Am. J. Med. Genet. A. 140A, 1219–1222.

[38] Õunap, K., Ilus, T. & Bartsch, O. (2005). A girl with inverted triplication of chromosome 3q25.3 → q29 and multiple congenital anomalies consistent with 3q duplication syndrome. Am. J. Med. Genet. A 134A, 434–438.

[39] Wu, C. et al. (2005). Mitochondrial DNA mutations and mitochondrial DNA depletion in gastric cancer. Genes Chromosomes Cancer 44, 19–28.

[40] Carvalho, C. M. B. et al. (2011). Inverted genomic segments and complex triplication rearrangements are mediated by inverted repeats in the human genome. Nat. Genet. 43, 1074.

[41] Brewer, B. J., Payen, C., Raghuraman, M. K. & Dunham, M. J. (2011). Origin-dependent inverted-repeat amplification: A replication-based model for generating palindromic amplicons. PLOS Genet. 7, e1002016.

[42] Martin, R. et al. (2024). Template switching between the leading and lagging strands at replication forks generates inverted copy number variants through hairpin-capped extrachromosomal DNA. PLOS Genet. 20, e1010850.

[43] Ho, S. S., Urban, A. E. & Mills, R. E. (2020). Structural variation in the sequencing era. Nat. Rev. Genet. 21, 171–189.

[44] Kosugi, S. & Terao, C. (2024). Comparative evaluation of SNVs, indels, and structural variations detected with short- and long-read sequencing data. Hum. Genome Var. 11, 18.

[45] Bolognini, D. et al. (2020). VISOR: a versatile haplotype-aware structural variant simulator for short- and long-read sequencing. Bioinformatics 36, 1267–1269.

[46] Dierckxsens, N., Li, T., Vermeesch, J. R. & Xie, Z. (2021). A benchmark of structural variation detection by long reads through a realistic simulated model. Genome Biol. 22, 342.

[47] Sedlazeck, F. J. et al. (2018). Accurate detection of complex structural variations using single-molecule sequencing. Nat. Methods 15, 461–468.

[48] Heller, D. & Vingron, M. (2019). SVIM: structural variant identification using mapped long reads. Bioinformatics 35, 2907–2915.

[49] Liu, Y., Kao, H.-I. & Bambara, R. A. (2004). Flap endonuclease 1: A central component of DNA metabolism. Annu. Rev. Biochem. 73, 589–615.

[50] Sun, H. et al. (2023). Okazaki fragment maturation: DNA flap dynamics for cell proliferation and survival. Trends in Cell Biology 33, 221–234.

[51] Sun, H. et al. (2021). Error-prone, stress-induced 3′ flap–based Okazaki fragment maturation supports cell survival. Science 374, 1252–1258.

[52] Winzeler, E. A. et al. (1999). Functional characterization of the *S. cerevisiae* genome by gene deletion and parallel analysis. Science 285, 901–906.

[53] Giaever, G. et al. (2002). Functional profiling of the *Saccharomyces cerevisiae* genome. Nature 418, 387–391.

[54] Wach, A., Brachat, A., Pöhlmann, R. & Philippsen, P. (1994). New heterologous modules for classical or PCR-based gene disruptions in *Saccharomyces cerevisiae*. Yeast 10, 1793–1808.

[55] Smith, D. J. & Whitehouse, I. (2012). Intrinsic coupling of lagging-strand synthesis to chromatin assembly. Nature 483, 434–438.

[56] Jiang, C. & Pugh, B. F. (2009). Nucleosome positioning and gene regulation: advances through genomics. Nat. Rev. Genet. 10, 161–172.

[57] Yuan, G.-C. et al. (2005). Genome-scale identification of nucleosome positions in *S. cerevisiae*. Science 309, 626–630.

[58] Mavrich, T. N. et al. (2008). Nucleosome organization in the Drosophila genome. Nature 453, 358–362.

[59] Lee, W. et al. (2007). A high-resolution atlas of nucleosome occupancy in yeast. Nat. Genet. 39, 1235–1244.

[60] Schones, D. E. et al. (2008). Dynamic regulation of nucleosome positioning in the human genome. Cell 132, 887–898.

[61] Connelly, J. C., Kirkham, L. A. & Leach, D. R. F. (1998). The SbcCD nuclease of *Escherichia coli* is a structural maintenance of chromosomes (SMC) family protein that cleaves hairpin DNA. Proc. Natl. Acad. Sci. U S A 95, 7969–7974.

[62] Dennenmoser, S. et al. (2017). Copy number increases of transposable elements and protein-coding genes in an invasive fish of hybrid origin. Mol. Ecol. 26, 4712–4724.

[63] Zichner, T. et al. (2013). Impact of genomic structural variation in drosophila melanogaster based on population-scale sequencing. Genome Res. 23, 568–579.

[64] Jeffares, D. C. et al. (2017). Transient structural variations have strong effects on quantitative traits and reproductive isolation in fission yeast. Nat. Commun. 8, 14061.

[65] Weischenfeldt, J., Symmons, O., Spitz, F. & O. Korbel, J. (2013). Phenotypic impact of genomic structural variation: insights from and for human disease. Nat. Rev. Genet. 14, 125–138.

[66] Chaisson, M. J. P., Wilson, R. K. & Eichler, E. E. (2015). Genetic variation and the de novo assembly of human genomes. Nat. Rev. Genet. 16, 627–640.

[67] Gong, L. et al. (2018). Picky comprehensively detects high resolution structural variants in nanopore long reads. Nat. Methods 15, 455–460.

[68] Schuler, B. A. et al. (2022). Lessons learned: next-generation sequencing applied to undiagnosed genetic diseases. J. Clin. Invest. 132, e154942.

[69] The 1000 Genomes Project Consortium (2015). A global reference for human genetic variation. Nature 526, 68–74.

[70] Chaisson, M. J. P. et al. (2019). Multi-platform discovery of haplotype-resolved structural variation in human genomes. Nat. Commun. 10, 1784.

[71] Chen, Y. et al. (2023). Deciphering the exact breakpoints of structural variations using long sequencing reads with DeBreak. Nat. Commun. 14, 283.

[72] Viguera, E., Canceill, D. & Ehrlich, S. D. (2001). Replication slippage involves DNA polymerase pausing and dissociation. EMBO J. 20, 2587–2595.

[73] Ohshima, K. & Wells, R. D. (1997). Hairpin formation during DNA synthesis primer realignment *in vitro* in triplet repeat sequences from human hereditary disease genes. J. Biol. Chem. 272, 16798–16806.

[74] Canceill, D., Viguera, E. & Ehrlich, S. D. (1999). Replication slippage of different DNA polymerases is inversely related to their strand displacement efficiency. J. Biol. Chem. 274, 27481–27490.

[75] Yarranton, G. T. & Gefter, M. L. (1979). Enzyme-catalyzed DNA unwinding: Studies on *Escherichia coli rep* protein. Proc. Natl. Acad. Sci. U.S.A. 76, 1658–1662.

[76] Heller, R. C. & Marians, K. J. (2007). Non-replicative helicases at the replication fork. DNA Repair (Amst) 6, 945–952.

[77] Cheng, W., Hsieh, J., Brendza, K. M. & Lohman, T. M. (2001). *E. coli* Rep oligomers are required to initiate DNA unwinding in vitro. J. Mol. Biol. 310, 327–350.

[78] Langmead, B. & Salzberg, S. L. (2012). Fast gapped-read alignment with Bowtie 2. Nat. Methods 9, 357–359.

[79] Camacho, C. et al. (2009). BLAST+: architecture and applications. BMC Bioinformatics 10, 421.

[80] Robinson, J. T. et al. (2011). Integrative genomics viewer. Nat. Biotechnol. 29, 24–26.

[81] Koboldt, D. C. et al. (2012). VarScan 2: Somatic mutation and copy number alteration discovery in cancer by exome sequencing. Genome Res. 22, 568–576.

[82] Wang, K., Li, M. & Hakonarson, H. (2010). ANNOVAR: functional annotation of genetic variants from high-throughput sequencing data. Nucleic Acids Res. 38, e164.

[83] Morgulis, A. et al. (2008). Database indexing for production MegaBLAST searches. Bioinformatics 24, 1757–1764.

[84] Chen, Y., Ye, W., Zhang, Y. & Xu, Y. (2015). High speed BLASTN: an accelerated MegaBLAST search tool. Nucleic Acids Res. 43, 7762–7768.

[85] Infante, J. J., Law, G. L. & Young, E. T. (2012). Analysis of nucleosome positioning using a nucleosome-scanning assay. In Chromatin Remodeling: Methods and Protocols (ed. Morse, R. H.) 63–87. Humana Press.

[86] Chen, K. et al. (2013). DANPOS: Dynamic analysis of nucleosome position and occupancy by sequencing. Genome Res. 23, 341–351.

